# Protective coding variants in *CFH* and *PELI3* and a variant near *CTRB1* are associated with age-related macular degeneration

**DOI:** 10.1101/034173

**Authors:** Erin K. Wagner, Yi Yu, Eric H. Souied, Sanna Seitsonen, Ilkka J. Immonen, Paavo Häppölä, Soumya Raychaudhuri, Mark J. Daly, Johanna M. Seddon

## Abstract

Although >20 common frequency age-related macular degeneration (AMD) alleles have been discovered with genome-wide association studies, substantial disease heritability remains unexplained. In this study we sought to identify additional variants, both common and rare, that have an association with advanced AMD. We genotyped 4,332 cases and 25,268 controls of European ancestry from three different populations using the Illumina Infinium HumanExome BeadChip. We performed meta-analyses to identify associations with common variants and performed single variant and gene-based burden tests to identify associations with rare variants. Two protective, low frequency, non-synonymous variants A307V in *PELI3* (odds ratio [OR]=0.14, *P*=4.3×10^−10^) and N1050Y in *CFH* (OR=0.76, *P*_conditional_=1.6×10^−11^) were significantly associated with a decrease in risk of AMD. Additionally, we identified an enrichment of protective alleles in *PELI3* using a burden test (OR=0.14). The new variants have a large effect size, similar to rare mutations we reported previously in a targeted sequencing study, which remain significant in this analysis: *CFH* R1210C (OR=18.82, *P*=3.5×10^−07^), *C3* K155Q (OR=3.27, *P*=1.5×10^−10^), and *C9* P167S (OR=2.04, *P*=2.8×10^−07^). We also identified a strong protective signal for a common variant (rs8056814) near *CTRB1* associated with a decrease in AMD risk (logistic regression: OR = 0.71, P = 1.8x10^−07^; Firth corrected OR = 0.64, P = 9.6x10^−11^). This study supports the involvement of both common and low frequency protective variants in AMD. It also may expand the role of the high-density lipoprotein pathway and branches of the innate immune pathway, outside that of the complement system, in the etiology of AMD.

## INTRODUCTION

Advanced age-related macular degeneration (AMD) (MIM 603075) is a common, complex, chronic eye disease (1). As a leading cause of vision loss in people older than 60 years, AMD currently affects more than 1.75 million individuals in the United States. This number is expected to increase by 50% to 3 million in 2020 due to aging of the population (2). The prevalence of AMD is expanding as the population ages, and therefore, the personal, societal, and economic burden is rising. The sibling recurrence-risk ratio (λ_s_) for AMD is estimated to be 3-6, suggesting that the risk of AMD is heavily influenced by genetic components, and twin studies have estimated the heritability of advanced AMD to be as high as 0.71 (3). Common variants in several alternative complement pathway genes, including complement factor H (*CFH*) (4–9), complement component 2 (*C2*)(8, 10), complement factor B (*CFB*)(8, 10), complement component 3 (*C3*) (11), and complement factor I (*CFI*) (12) and a variant in the age-related maculopathy susceptibility 2 (*ARMS2*) gene (13, 14) modulate AMD risk. Genome-wide association studies (GWAS) in large cohorts have also identified common variants in the high-density lipoprotein cholesterol (HDL), extracellular collagen matrix and angiogenesis pathways (15–17). A metaanalysis confirmed the above loci and added new loci for AMD through GWAS and extensive imputation approaches, yielding a total of 19 significant associations in common loci (18).

Despite a rapidly growing list of associations with common variants, there continues to be a large proportion of the heritability of AMD that is unexplained (17, 18), which might be due to undiscovered common variants or rare alleles in the genome. In many instances linking the associated variant to causal risk-conferring functional variation has been challenging (19). Since common variants have survived the effects of purifying negative selection, they often, by necessity, have subtle biochemical or regulatory functions that can be difficult to assess functionally (20). Sequencing can open up the whole spectrum of the allele frequency distribution and detect rare mutations with obvious functional consequences. In fact, recent studies using sequencing approaches have successfully identified several rare functional variants in *CFH, C3, CFI* and complement component 9 (*C9*) that may have direct impact on the activation of alternative complement cascade (21–25). A cost-effective alternative approach to query the functional variants across the whole exome is to use an exome array, which provides good coverage for functional variants with frequency as low as 0.01%. To examine the spectrum of rare variation in the exome, we genotyped large cohorts of individuals of European ancestry using the Illumina Infinium HumanExome BeadChip with custom content of loci related to AMD (3,214 additional custom variants).

## RESULTS

### Variants passing quality control

We genotyped all samples using the Illumina Infinium HumanExome BeadChip with custom content, of which 161,374 (64.2%) variants are polymorphic in our samples and passed quality control. We then categorized 40,087 variants as common (minor allele frequency [MAF] ≥ 1%), and 121,287 variants as rare (MAF < 1%) based on their minor allele frequency in cases and in controls. This platform captured 1.7% (121,287/7,330,859=.017) of possible rare variant sites in the exome as compared to the >7,300,000 rare variant sties identified in release 0.3 of the Exome Aggregation Consortium (ExAC) (26). Among those rare variants, 72,503 variants were nonsynonymous, nonsense or splice-site variants.

### Rare variant analysis

We first tested for association with the rare variants included on the Illumina Infinium HumanExome BeadChip with custom content. After strict quality control and filtering, a total of 57,101 variants were tested for association with AMD. A low frequency, nonsynonymous variant in the pellino E3 ubiquitin protein ligase family member 3 gene (*PELI3*), rs145732233 (*PELI3* A307V), was significantly associated at the genome-wide level with AMD (odds ratio [OR] = 0.14, *P* = 4.3 × 10^−10^) (**Table 1**). This variant was predicted to be ‘possibly damaging’ by PolyPhen2. In our samples, rs145732233 has a minor allele frequency of 0.21% in cases versus 0.77% in controls (data not shown). Comparatively, the MAF is 0.54% in the ExAC database, which is lower than its frequency in our control group and higher than its frequency in our case group. This indicates that the protective effect of A307V in the *PELI3* gene is likely to be true, although the effect size might be smaller than the value estimated in our samples. Another rare, nonsynonymous SNP in *CFH*, rs35274867 (*CFH*N1050Y), was also significantly associated at a genome-wide level with AMD (OR = 0.76, *P* = 6.2x10^−12^). We tested this SNP for independence of the other known common and rare SNPs in the *CFH* gene and found the association signal did decrease slightly in the conditional analysis, but the association signal was still highly significant (*P* = 1.6x10^−11^). We also detected several other loci with suggestive evidence of association (1x10^−05^ > *P* > 5x10^−08^) with AMD (**Supplementary Table S2**).

**Table 1.**
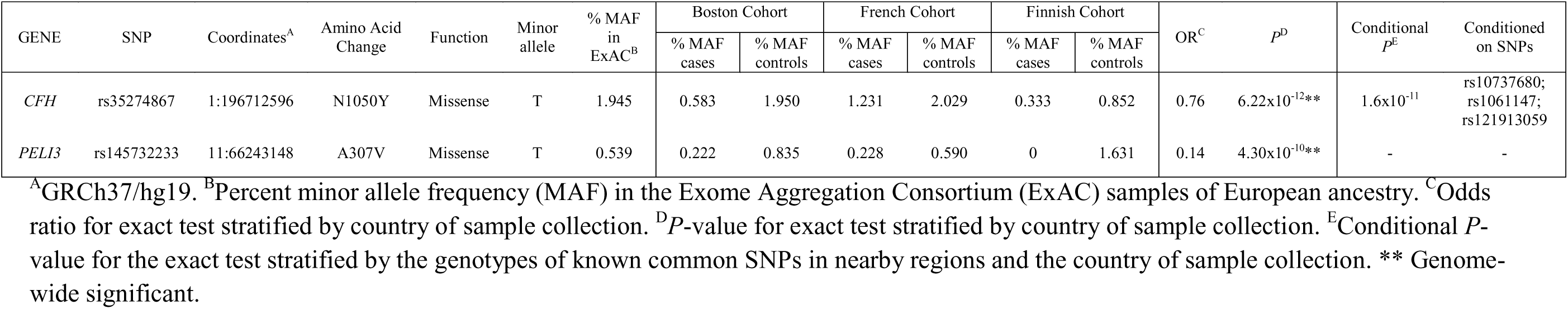
New rare or low frequency age-related macular degeneration associated variants: associations in the single-SNP analysis.

| GENE | SNP | Coordinates<sup>A</sup> | Amino Acid Change | Function | Minor allele | % MAF in ExAC<sup>B</sup> | Boston Cohort |  | French Cohort |  | Finnish Cohort |  | OR<sup>C</sup> | *P*<sup>D</sup> | Conditional *P*<sup>E</sup> | Conditioned on SNPs |
| --- | --- | --- | --- | --- | --- | --- | --- | --- | --- | --- | --- | --- | --- | --- | --- | --- |
|  |  |  |  |  |  |  | % MAF cases | % MAF controls | % MAF cases | % MAF controls | % MAF cases | % MAF controls |  |  |  |  |
| *CFH* | rs35274867 | 1:196712596 | N1050Y | Missense | T | 1.945 | 0.583 | 1.950 | 1.231 | 2.029 | 0.333 | 0.852 | 0.76 | 6.22x10<sup>-12</sup>** | 1.6x10<sup>-11</sup> | rs10737680;<br>rs1061147;<br>rs121913059 |
| *PELI3* | rs145732233 | 11:66243148 | A307V | Missense | T | 0.539 | 0.222 | 0.835 | 0.228 | 0.590 | 0 | 1.631 | 0.14 | 4.30x10<sup>-10</sup>** | - | - |

<sup>A</sup>GRCh37/hg19. <sup>B</sup>Percent minor allele frequency (MAF) in the Exome Aggregation Consortium (ExAC) samples of European ancestry. <sup>C</sup>Odds ratio for exact test stratified by country of sample collection. <sup>D</sup>*P*-value for exact test stratified by country of sample collection. <sup>E</sup>Conditional *P*-value for the exact test stratified by the genotypes of known common SNPs in nearby regions and the country of sample collection. \*\* Genome-wide significant.

In addition to testing each single-variant individually, we also performed gene-based tests to further investigate the cumulative effects of rare functional variants in AMD. Even though the distribution of association statistics of gene-based tests were slightly deflated, possibly due to lack of power from genes with small numbers of rare variants, we were still able to detect significant signals in several genes (inflation factor λ_gc_ = 0.96, **Supplementary S1**). We identified a burden of rare variants in the *PELI3* gene (*P* = 4.3x10^−07^) using the simple burden analysis. We also found cumulative effects of rare variants in five other genes to be significantly associated at the genome-wide level with AMD: *CFH*, *C3*, *C9*, abnormal spindle microtubule assembly (*ASPM*), and mutS homolog 5 (*MSH5*) using either the simple burden test or the SKAT analysis (**Table 2**). To test if the significant signals were independent of known variants, we conditioned on nearby known common and rare variants and demonstrated that the signals were mostly driven by the known variants.

**Table 2.** Gene based analysis for burden of rare variants in age-related macular degeneration.

| Gene | Chromosome | Number of variants<sup>A</sup> | $P_{\text{Burden}}^{\text{B}}$ | $P_{\text{SKAT}}^{\text{C}}$ | Conditional $P_{\text{SKAT}}^{\text{D}}$ | Conditioned on variants |
| --- | --- | --- | --- | --- | --- | --- |
| *PELI3* | 11 | 3 | $4.3 \times 10^{-07*}$ | $4.3 \times 10^{-06}$ | $5.5 \times 10^{-01}$ | rs145732233 |
| *MSH5* | 6 | 7 | $2.4 \times 10^{-08*}$ | $1.1 \times 10^{-07*}$ | $1.5 \times 10^{-01}$ | rs429608; rs3129987 |
| *C9* | 5 | 13 | $1.1 \times 10^{-04}$ | $9.8 \times 10^{-08*}$ | $3.0 \times 10^{-01}$ | rs34882957 |
| *CFH* | 1 | 12 | $3.8 \times 10^{-12*}$ | $2.8 \times 10^{-15*}$ | $5.1 \times 10^{-01}$ | rs1061147; rs10737680; rs121913059; rs35274867 |
| *ASPM* | 1 | 31 | $1.1 \times 10^{-05}$ | $3.5 \times 10^{-07*}$ | $7.3 \times 10^{-01}$ | rs1061147; rs10737680; rs121913059; rs35274867 |
| *C3* | 19 | 10 | $1.1 \times 10^{-03}$ | $1.7 \times 10^{-12*}$ | $7.9 \times 10^{-01}$ | rs2230199; rs147859257 |

\*Genes with $P < 2.06 \times 10^{-06}$ . <sup>A</sup>Number of rare variants passing stringent quality control in each gene used for each test. <sup>B</sup> $P$ -value of gene-based simple burden analysis. <sup>C</sup> $P$ -value of gene-based SKAT analysis. <sup>D</sup> $P$ -value of the conditional gene-based SKAT analysis for each gene adjusting for the genotypes of common and rare variants in and near the gene.

We additionally examined the association signals at previously reported rare AMD loci in *CFH*, *CFI*, *C3* and *C9* in our recent targeted sequencing study, from which there are a subset of samples that overlap with those used for this genotyping study, and found the results supported those obtained from the prior analysis (21, 22) (**Supplementary Table S1**).

### Common variant analysis

We evaluated common variants using logistic regression, adjusting for genetic ancestry based on principal component analysis, analyzing the samples from the Boston, French and Finnish cohorts separately and then performed a meta-analysis to assess the pooled effect of these variants across the three countries. We plotted the *P*-values of 15,671 ancestry-informative markers in Quantile-Quantile plots and observed no statistical inflation in the distribution of the association statistic for any of the three country-of-origin specific logistic regression analyses (genomic inflation factor λ_gc_ = 1.0).

**Table 3** shows the newly associated, common variants identified in the advanced AMD analysis. We identified one independent significant signal associated with advanced AMD on chromosome 16 near the chymotrypsinogen B1 gene (*CTRB1*) (logistic regression: OR = 0.71, *P* = 1.8x10^−07^; Firth corrected OR = 0.64, *P* = 9.6x10^−11^; **Figure 1**) which is not close to any of the known loci. The top variant in this region, rs8056814, is an intergenic single nucleotide polymorphism upstream from *CTBR1.*

**Table 3.** New common age-related macular degeneration associated variants: meta-analysis of age-related macular degeneration cohorts.

| Gene | SNP | Coordinates<sup>A</sup> | Minor allele | Boston cohort |  |  |  | French cohort |  |  |  | Finnish cohort |  |  |  | Meta-analysis |  |  | Firth Corrected Meta-analysis |  |
| --- | --- | --- | --- | --- | --- | --- | --- | --- | --- | --- | --- | --- | --- | --- | --- | --- | --- | --- | --- | --- |
|  |  |  |  | MAF cases | MAF controls | OR | P | MAF cases | MAF controls | OR | P | MAF cases | MAF controls | OR | P | OR<sup>B</sup> | P<sup>C</sup> | Direction<sup>D</sup> | OR<sup>E</sup> | P<sup>F</sup> |
| *CTRB1* | rs8056814 | 16:75252327 | A | 0.072 | 0.104 | 0.67 | 2.5x10<sup>-07</sup> | 0.080 | 0.088 | 0.90 | 5.1x10<sup>-01</sup> | 0.063 | 0.083 | 0.75 | 9.1x10<sup>-02</sup> | 0.71 | 1.8x10<sup>-07*</sup> | --- | 0.64 | 9.6 x10<sup>-11</sup> |
| *APOH* | rs1801689 | 17:64210580 | C | 0.028 | 0.038 | 0.71 | 1.5x10<sup>-03</sup> | 0.029 | 0.048 | 0.56 | 3.6x10<sup>-04</sup> | 0.007 | 0.008 | 0.83 | 7.2x10<sup>-01</sup> | 0.67 | 4.1x10<sup>-06*</sup> | --- | 0.74 | 7.7 x10<sup>-04</sup> |
| *COL4A3* | rs11884770 | 2:228086920 | T | 0.251 | 0.280 | 0.85 | 1.4x10<sup>-03</sup> | 0.248 | 0.315 | 0.72 | 1.3x10<sup>-03</sup> | 0.308 | 0.337 | 0.87 | 1.4x10<sup>-01</sup> | 0.83 | 6.6x10<sup>-06*</sup> | --- | 0.88 | 4.0 x10<sup>-03</sup> |
| *MUC1* | rs4072037 | 1:155162067 | C | 0.453 | 0.476 | 0.89 | 1.6x10<sup>-03</sup> | 0.444 | 0.495 | 0.80 | 5.8x10<sup>-04</sup> | 0.423 | 0.438 | 0.94 | 4.6x10<sup>-01</sup> | 0.88 | 9.6x10<sup>-06*</sup> | --- | 0.91 | 1.9 x10<sup>-03</sup> |

Common variants (minor allele frequency [MAF] $\geq 1\%$ ) in cases and in controls across all three populations with significant ( $P$ -value $< 1.24 \times 10^{-06}$ ) or suggestive ( $P < 2.49 \times 10^{-05}$ ) associations with advanced AMD as discovered in the meta-analysis. <sup>A</sup>GRCh37/hg19. <sup>B</sup>Odds ratio from the meta-analysis of logistic regression analysis of common variants. <sup>C</sup> $P$ -value from the meta-analysis of logistic regression analysis of common variants. <sup>D</sup>Direction of effect from the three populations in the following order: Boston, French, Finnish. <sup>E</sup>Odds ratio from the meta-analysis of Firth logistic regression analysis of common variants. <sup>F</sup> $P$ -value from the meta-analysis of Firth logistic regression analysis of common variants. \*Genome-wide suggestive.

**Figure 1.**
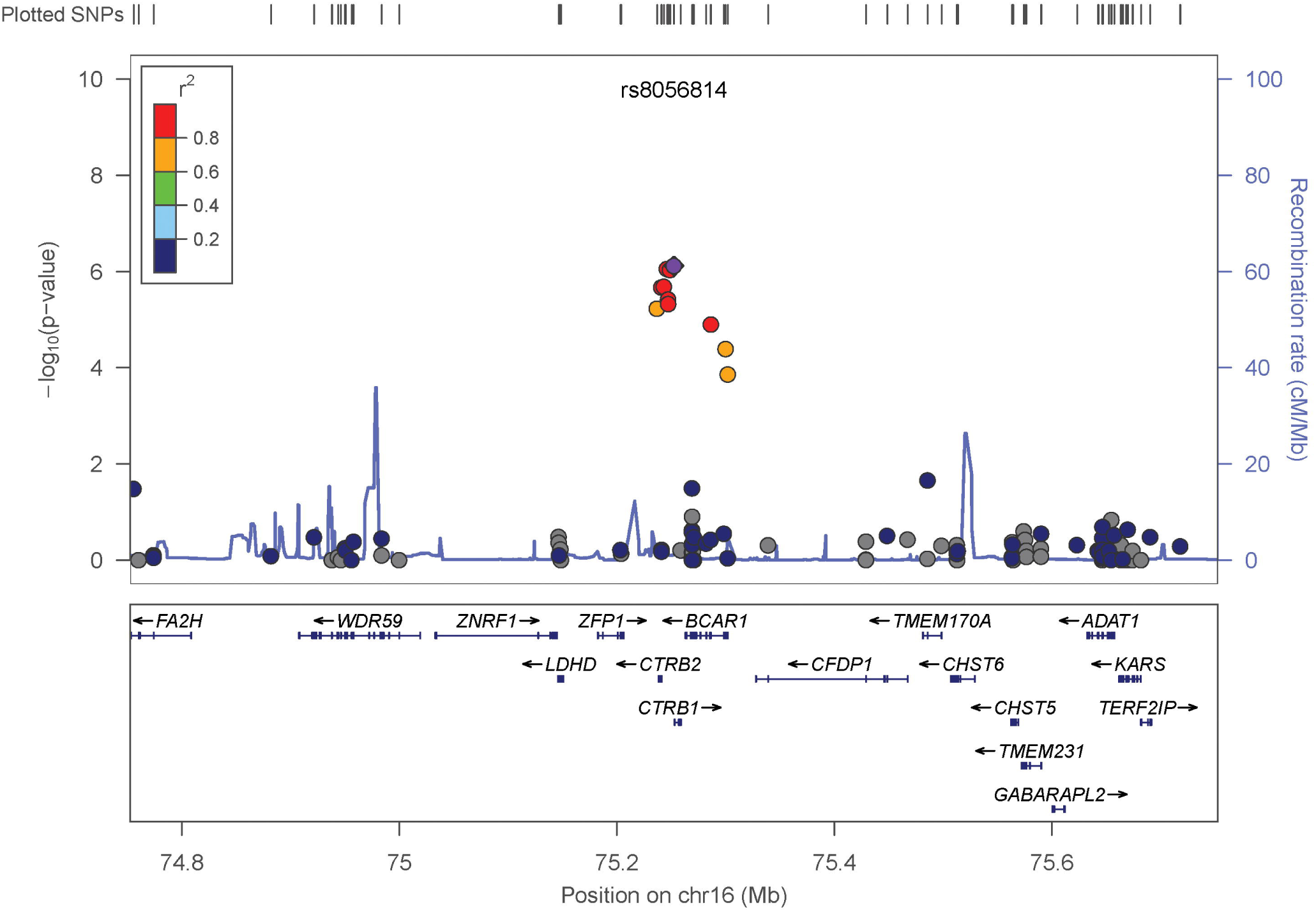
Zoomplots summarizing association results for the CTRB1 locus. The regional association plot from analysis of common variants in 4,332 cases and 4,642 controls of European ancestry. Gene location is shown along the bottom of the graph, with observed -log10(*P*-value) along the left Y-axis and recombination rate along the right Y-axis. Each variant is plotted as a circle, filled with color coded according to the extent of linkage disequilibrium (variants with missing LD information are shown in grey) with the index variant, rs8056814 (colored in purple).

We also identified three additional suggestive common variant associations with advanced AMD (**Table 3**). These variants included: a missense variant, rs1801689 in apolipoprotein H (*APOH*) (C325R); an intronic variant, rs11884770 in collagen, type IV, alpha 3 (*COL4A3*); and a synonymous variant, rs4072037 in mucin 1, cell surface associated (*MUC1*). We identified seven variants with a borderline suggestive association with AMD, including variants in genes involved in the complement and the innate immune system pathways, two pathways known to be involved in the etiology of AMD (**Supplementary Table S2**). **Supplementary Table S3** shows common variants or their proxy at 21 loci in the inflammatory/immune, angiogenesis, collagen/extracellular matrix, and lipid pathways previously identified by several GWAS analyses (4, 15–18). All known alleles showed similar effect size and direction in this study as compared to previously published values.

For the AMD subtype analyses (**Supplementary Table S4**), we found no new genome-wide significant associations between the two advanced subtypes, geographic atrophy and neovascular disease, nor did we find any new genome-wide significant association between either of the subtypes and the control group. We did, however, identify a genome-wide suggestive association between geographic atrophy and the missense variant, rs1715828 in dynein assembly factor with WDR repeat domains 1 (*DAW1*) (T121S) (*P* = 5.5x10^−06^). Genome-wide suggestive associations with neovascular disease include: rs7604613, an intragenic variant near tetratricopeptide repeat domain 32 (*TTC32*) (*P* = 1.7x10^−05^) and both rs11884770 (*P* = 1.7x10^−05^) and rs1801689 (*P* = 1.9x10^−06^), the same variants found to be suggestively associated with advanced AMD. The missense variant rs1801689 in *APOH* shows a stronger association with neovascular disease than with advanced AMD (OR_neovascular_ = 0.61 vs. OR_overall_advanced_ = 0.67), while rs11884770 in *COL4A3* shows a weaker association with neovascular disease than with overall combined types of advanced AMD (OR_neovascular_ = 0.81 vs. OR_overall_advanced_ = 0.83). Confirming known genome-wide associations with AMD, we identified genome-wide significant associations between geographic atrophy and the known loci including *CFH*, *C3*, *C2*/*CFB*, and *ARMS2* and between neovascular disease and known AMD loci including *CFH*, *COL8A1*, *C2*/*CFB*, *C9*, *TGFBR1*, *CETP*, *C3*, and *TIMP3.* The association analysis testing the difference between the two subtypes identified and confirmed the known association with *ARMS2* as previously reported (27, 28), but did not identify any novel associations with any other variants.

## DISCUSSION

In this study, we aimed to find new genetic factors for advanced AMD by querying common and rare functional variants across the exome in a large number of subjects from three cohorts of European ancestry. We identified significant associations between AMD and a rare or low frequency protective missense variants, *PELI3* A307V (*P* = 4.3×10^−10^) and *CFH*N1050Y (*P* = 6.2x10^−12^) and a suggestively associated common protective variant near *CTRB1* (rs8056814, *P* = 1.8x10^−07^).

The rare non-synonymous variant, rs145732233 in *PELI3* is newly identified in this study and is associated with a decrease in risk of AMD. A burden of rare variants in *PELI3* was also detected in the simple burden test, but just missed the cutoff for significance in the sequence kernel association test (SKAT) analysis. The simple burden test also shows the overall association signal from *PELI3* is protective (OR = 0.28, data not shown). *PELI3* encodes E3 ubiquitin ligase pellino, a scaffold protein that helps transmit the immune response signals. Pellino E3 has been demonstrated to augment the expression of type I interferon but not of proinflammatory cytokines in response to toll-like receptor 3 protein (TLR3) activation (29). Similar to the complement pathway, the Toll-like receptors (TLRs) participate in protective response against microbial invasion when activated normally, but could be harmful to the host when activated improperly or uncontrolled (30). A protective variant such as *PELI3* A307V might enhance the signaling transduction between the TLR3 and IRF7 pathways, resulting in downregulation of type I interferon expression (29). Thus, individuals with the *PELI3* A307V mutation may have less severe type I interferon response than individuals with normal pellino E3. Individuals with the *PELI3* A307V mutation may be protected from damage caused by immune response activated abnormally, and therefore could be less likely to develop advanced AMD. Although variants in *TLR3* (31) and toll-like receptor 3 (*TLR4*) (32) have been suspected to be related to AMD, these common associations were not replicated in studies with large cohorts (15–18). By genotyping a large number of samples using the exome array, this study identified a novel rare mutation in a gene that intermediates signals between Toll-like receptors and the innate immune pathway. Further studies of functional roles of this mutation and possible mechanisms associated with AMD are warranted, but considering the results from the rare variant analyses, variants in *PELI3* may play a protective role in AMD.

Rare variant analysis in this study identified a second protective rare variant significantly associated with a decrease in risk of AMD. The missense SNP, rs35274867 (*CFH*N1050Y) (OR = 0.76; *P* = 6.2x10^−12^) is located in *CFH*, a gene in the complement pathway with strong associations with risk of AMD. An association analysis of the *CFH*N1060Y variant conditional on the associated risk variants in *CFH* shows this SNP has an independent effect on AMD (**Table 1**). Haplotype construction using the Boston cohort also shows the protective allele at *CFH* N1050Y occurs independently of the risk variants from *CFH* R1210C (**Table 4**). Other studies have reported an association between *CFH* variants, both common and rare, and AMD (4–9, 22, 33, 34). This is the first time *CFH*N1050Y has been significantly and independently associated at the genome-wide level with AMD. This variant was identified in our previous sequencing studies, but was not independently associated at the genome-wide level with AMD at the time, though the variant was noted as having a higher frequency in controls than in AMD cases (22, 25). Additionally, the variant has previously been found with a higher frequency in controls compared to cases in a study of systemic lupus erythematosus (35), and was found at a higher frequency in controls than cases in a study of atypical hemolytic uremic syndrome, a disease that shares genetic influences with AMD. Conversely, there is evidence of affected patients carrying the variants including a father and daughter with basal laminar drusen (36), though this study did not look at the frequency of the variant in controls, nor did it attribute the variant to causing the disease in these two patients, and two patients with hemolytic uremic syndrome (37). The variant is located in complement control protein module 18 (CCP18), a module that harbors multiple disease-associated loci and may to be part of an essential binding site (38). Alteration of binding domains of *CFH* may modify how the factor H protein regulates the alternative complement pathway. As this variant is associated with a decrease in risk of AMD, it suggests the mutation may create a comparatively enhanced binding site. Further study is needed to determine the underlying biological mechanism and protective function of this variant.

**Table 4.**
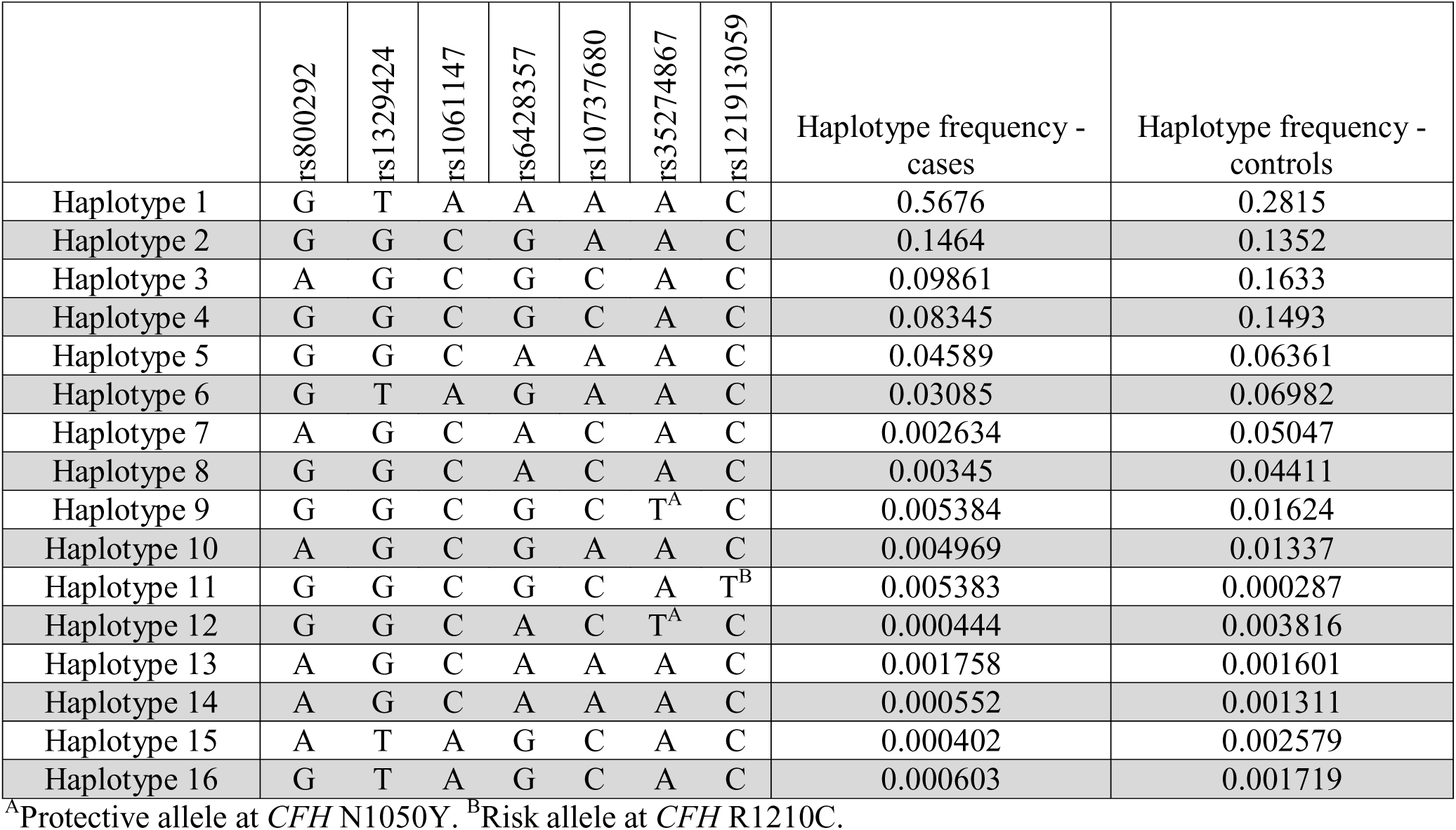
CFH Haplotype construction in the Boston cohort. Protective allele at *CFH* N1050Y. Risk allele at *CFH* R1210C.

|  | rs800292 | rs1329424 | rs1061147 | rs6428357 | rs10737680 | rs35274867 | rs121913059 | Haplotype frequency -<br>cases | Haplotype frequency -<br>controls |
| --- | --- | --- | --- | --- | --- | --- | --- | --- | --- |
| Haplotype 1 | G | T | A | A | A | A | C | 0.5676 | 0.2815 |
| Haplotype 2 | G | G | C | G | A | A | C | 0.1464 | 0.1352 |
| Haplotype 3 | A | G | C | G | C | A | C | 0.09861 | 0.1633 |
| Haplotype 4 | G | G | C | G | C | A | C | 0.08345 | 0.1493 |
| Haplotype 5 | G | G | C | A | A | A | C | 0.04589 | 0.06361 |
| Haplotype 6 | G | T | A | G | A | A | C | 0.03085 | 0.06982 |
| Haplotype 7 | A | G | C | A | C | A | C | 0.002634 | 0.05047 |
| Haplotype 8 | G | G | C | A | C | A | C | 0.00345 | 0.04411 |
| Haplotype 9 | G | G | C | G | C | T<sup>A</sup> | C | 0.005384 | 0.01624 |
| Haplotype 10 | A | G | C | G | A | A | C | 0.004969 | 0.01337 |
| Haplotype 11 | G | G | C | G | C | A | T<sup>B</sup> | 0.005383 | 0.000287 |
| Haplotype 12 | G | G | C | A | C | T<sup>A</sup> | C | 0.000444 | 0.003816 |
| Haplotype 13 | A | G | C | A | A | A | C | 0.001758 | 0.001601 |
| Haplotype 14 | A | G | C | A | A | A | C | 0.000552 | 0.001311 |
| Haplotype 15 | A | T | A | G | C | A | C | 0.000402 | 0.002579 |
| Haplotype 16 | G | T | A | G | C | A | C | 0.000603 | 0.001719 |

<sup>A</sup>Protective allele at *CFH* N1050Y. <sup>B</sup>Risk allele at *CFH* R1210C.

The burden analyses detected a suggestive association signal (*P*_Burden_ = 3.2x10^−05^, *P*_SKAT_ = 1.0x10^−02^) in the *CFI* gene, but not as strong as the association signal we recently found in our sequencing study by simple burden tests (21). This may be due to the fact that there were only 12 *CFI* rare variants detected on the exome array, much less compared to the 59 *CFI* rare variants we detected by targeted sequencing. This exemplifies the value of employing a variety of genotyping and sequencing platforms for gene discovery.

We found an association between AMD and a common protective variant rs8056814 in *CTRB1.* This SNP is about 330kb away from rs8053796 in contactin associated protein-like 4 (*CNTNAP4*), where we detected a suggestive association signal (*P* = 1.7x10^−05^) in our previous meta-GWAS study (17). Previous studies have reported nominal associations between rs8056814 and AMD in the same direction of effect, and a recent, independent but concurrent study, reported a genome-wide significant association between rs8056814 and AMD (*P* = 2.9x10^−10^), also in the same direction of effect (18, 39–41). In our study, the association signal in the *CTRB1* locus reached statistical significance (*P* < 1.24x10^−06^) for the first time. Compared with previous studies, rs8056814 was directly genotyped in our analysis instead of being imputed, which enabled us to assess the association at this locus more accurately and with more power. Therefore, the effect of this locus might be underestimated in studies using imputed datasets as the linkage disequilibrium structure may not perfectly capture the genotype information of this locus.

The rs8056814 variant is located 557bp upstream of *CTRB1*, encoding chymotrypsinogen, a serine protease that is secreted into the gastrointestinal tract and activated by proteolytic cleavage with trypsin. *CTRB1* is a known risk locus for type 1 diabetes (42) and it is reported to be associated with variation in HDL levels as well (43). Querying the GTEx database, we found that rs8056814 is a cis-expression quantitative trait locus (eQTL) for the downstream gene, *BCAR1* in whole blood (*P* = 1.5x10^−07^, β = 0.37) (The data was obtained from the GTEx Portal and dbGaP accession number phs000424.v6.p1) (44). CHIPseq results from the ENCODE project suggests rs8056814 lies in the promoter region of *CTRB1* in a hepatic carcinoma cell line and in an enhancer region in blood cell lines which could explain the eQTL association with *BCAR1* in blood (45). Eye tissue was not assessed as part of the GTEx and ENCODE projects, so further work is required to determine the role this variant plays in ocular tissues. Further evidence shows another variant, rs7202877, is in high LD with rs8056814 (r^2^ = 0.9) and regulates expression levels of *CTRB1* (46). If *CTRB1* does play a role in the HDL pathway as suggested by Dastani et al. (43), the role of HDL pathway genes in AMD is not unprecedented. Variants in several HDL pathway genes, such as hepatic lipase (*LIPC*) (15, 16), plasma cholesteryl ester transfer protein (*CETP*) (15–17) and ATP binding cassette subfamily A member 1 (ABCA1) (15, 16, 47, 48) show significant associations with AMD. The apolipoprotein E (*APOE*) gene, involved in a different part of the lipid pathway, is also related to AMD (49). Of note, the minor allele for the associated variant in *LIPC* (rs10468017), which is associated with higher HDL levels, also shows a protective effect in AMD (15, 16). Considering the effects of other variants in this region, it is possible that the rare allele of rs8056814 could influence metabolic changes in serum glucose and HDL levels through the modulation of *CTRB1* expression levels.

The Illumina Infinium HumanExome BeadChip provides an alternative to whole exome sequencing as a way to assess functional variants across the exome in an unbiased manner. However, it does not provide complete coverage of all functional variants in all genes. The ultra-rare variants (MAF < 0.03%) are not covered by this array, thus limiting our power to detect individual or cumulative effects of these variants. For example, it includes only 12 rare variants in the *CFI* gene, while our recent targeted sequencing analysis detected 59 rare variants (21). The lack of information on those ultra-rare variants could weaken the association signals in the gene-based test, especially in the scenario when most of those ultra-rare variants in this gene are causal and associated with AMD in the same direction.

In summary, we have identified protective AMD loci in *CFH*, *PELI3* and near *CTRB1*. We provide new suggestive loci worthy of follow-up, including common variants in *COL4A3*, *APOH*, and *MUC1*, as well as several rare variants. We also confirmed previously published common loci in several pathways identified by GWAS, and the recently associated rare AMD loci in the complement genes *CFH*, *CFI*, *C3*, and *C9* discovered by targeted sequencing. The new genetic loci associated at the genome-wide level with AMD suggest that genes in other branches of the innate immune pathway and additional genes in the HDL pathway may also be involved in the etiology of AMD. As the knowledge about the genetic architecture of AMD expands, new variants may enhance predictive models (50, 51), and could lead to the development of new therapeutic targets.

## MATERIALS AND METHODS

### Case-control definitions

All individuals were evaluated by a board-certified ophthalmologist who conducted ocular examinations including with visual acuity measurements, dilated slit-lamp biomicroscopy, and stereoscopic color fundus photography. Ophthalmologic medical records and ocular images were also reviewed. All subjects were graded using the Clinical Age-Related Maculopathy Staging (CARMS) system (52). Case patients had either geographic atrophy (advanced central or non-central non-exudative AMD or CARMS grade 4) or neovascular disease (neovascular AMD or CARMS grade 5). Controls did not have early, intermediate or advanced macular degeneration, and were categorized as CARMS grade 1. All controls were 60 ≥ years old.

Boston cohorts were recruited at the Tufts Medical Center in Boston, Massachusetts, U.S.A., and throughout the country through ongoing AMD study protocols, as previously described (3, 8, 11, 12, 15, 53–55). We selected 3,772 unrelated individuals (2,488 case and 1,284 controls) from our large collection of advanced AMD case-control and family cohorts. The French cohort of 1,544 cases and 289 controls was recruited at Hôpital Intercommunal de Créteil, Créteil, France, as previously described (15, 17). The Finnish cohort of 300 cases and 160 controls was recruited at the Helsinki University Central Hospital, Helsinki, Finland and >20,000 additional Finnish controls were recruited as part of The National FINRISK Study, a cross-sectional survey of the Finnish population to assess chronic disease in people aged 25 to 74 (56, 57). We included genotype data for shared controls of 2,909 samples that had been genotyped at the Broad Institute (29, 58–61) (**Supplementary Table S5**). Individuals from all cohorts were self-reported white individuals of European descent. We used the first five principal components generated by EIGENSTRAT (62) based on the ancestry informative markers to calculate Euclidean distances between samples in the Boston and French cohorts and shared control samples. We then randomly selected individual case samples in these cohorts and assigned the nearest unassigned shared controls to the selected case’s cohort. We matched 2,434 of these shared controls to the Boston cohort and 475 of these shared controls to the French cohort. The two cohorts with matched controls were reexamined with five outlier removal iterations in EIGENSTRAT to ensure that samples were matched properly by their ethnic background.

### Whole exome array genotyping of coding variants

Genotyping was performed using the Illumina Infinium HumanExome BeadChip (v1.0), which provides coverage of over 240,000 functional exonic variants selected from >12,000 whole exome and known variants associated with complex traits in previous GWAS, human leukocyte antigen tags, ancestry-informative markers, markers for identity-by-descent estimation and random synonymous single nucleotide polymorphisms (SNPs) (http://genome.sph.umich.edu/wiki/Exome_Chip_Design). In addition, we customized our assay by adding 3,214 SNPs from candidate AMD genes and genes in associated pathways. Included in the custom content are common variants which achieved a *P*-value less than 0.001 in our previous meta-GWAS studies (15, 17) and 20 common SNPs reported by the AMDGENE consortium meta-GWAS study (18). We conducted genotyping of the Boston, French and Finnish cohort samples at the John Hopkins Genotyping Core Laboratory. We genotyped shared control samples separately at the Broad Institute using the same genotyping platform and custom content as was used for the Boston, French, and Finnish cohort samples. We called genotypes using Illumina’s GenomeStudio software and then used zCall (63), a rare-variant caller developed at the Broad Institute, to recover missed rare genotypes.

### Statistical Analyses

We required that samples have <2% missing genotype calls for common variants (MAF > 5%) before applying zCall. Then after applying zCall we removed duplicate variants, monomorphic variants, variants with a low call rate (<98%), and variants failing Hardy-Weinberg (P < 10^−06^). We merged genotype calls from the different cohorts by only including variants that passed quality control and passed the Hardy-Weinberg test (*P* ≥ 10^−06^) across all samples. To eliminate any batch effect, we excluded 412 variants with allele frequencies significantly different between the examined controls genotyped at the John Hopkins Core Laboratory and the shared controls genotyped at the Broad Institute (*P* < 10^−03^). We identified 15,671 ancestry informative markers with high minor allele frequencies (MAF > 5%), and excluded regions near (<1Mb) any of the 20 known AMD loci (18) and the major histocompatibility complex locus (chr 6, 25.0-35.0 Mb). We then pruned the resulting set of variants using the --indep option in PLINK with default parameters (variance inflation factor = 2, window size = 50 SNPs) (64). We assessed relatedness by calculating genome-wide proportion identity-by-descent estimates (PIHAT values) using these ancestry informative markers (64). We identified pairs of sequenced individuals with PIHAT > 0.2, and removed one of those individuals from the analysis. We then used EIGENSTRAT (62) to generate the first 10 principal components based on the ancestry informative markers. We only included shared controls who matched the genetic background of Boston and French samples based on principal components as described previously (21).

For statistical analysis of common variants, we tested for associations assuming an additive genetic model using logistic regression adjusting for the first 10 principle components from the EIGENSTRAT analysis. The summary data were then meta-analyzed using METAL (65). Firth logistic regression analysis, adjusted for the top 10 principal components was performed on the most significant associations using the logistf package in R. In addition to testing each variant for an association with advanced AMD, we also conducted subtype analyses and tested each variant for an association with the two advanced forms of AMD: neovascular disease and geographic atrophy. To recognize independent association signals, we also performed conditional analysis for the significant variants within 1Mb of any of the 20 known AMD loci (**Supplementary Table S3**) by adjusting for the genotype of the adjacent known variant. Genome-wide significance (*P*-value < 5×10^−08^) and genome-wide suggestive (*P*-value < 1×10^−05^) thresholds were used evaluate the association signals of common variants.

For rare functional variants we carried out single-variant association tests using the same statistical framework of exact statistics described previously (21, 22). Briefly, we used a 2×2 Fisher’s exact test to calculate a one-tailed exact *P*-value for multiple case-control cohorts. We performed analyses on 57,101 variants with a MAF <1%, and included. The first 10 principal components of EIGENSTRAT were included as covariates in this analysis. We applied the same statistical framework on data further stratified by genotypes of nearby known common variants, in addition to country of sample collection, to achieve the *P*-values of conditional analysis for the rare variants. To eliminate potential false positives due to low quality calls of rare variants, we also re-examined the cluster plots of genotype calls for the significant variants after association tests, and excluded variants poorly clustered. Genome-wide significance (*P*-value < 5×10^−08^) and genome-wide suggestive (*P*-value < 1×10^−05^) thresholds were used evaluate the association signals of common variants.

For gene-based analysis, we performed a SKAT analysis on each cohort separately followed by a metaanalysis implemented using the RAREMETAL software, to assess the pooled effect of the rare variant burden across the three cohorts (66). SKAT has been shown to perform well in scenarios when a large fraction of the variants in a region are non-causal or the effects of causal variants are in different directions. We performed analyses using default weights (67) on 72,503 nonsynonymous, nonsense or splice-site variants with MAF between 1 and 0.03% in cases or in controls groups. We excluded extremely rare variants (minor allele count < 5, or MAF < 0.03%) from this test. These variants are located in 12,122 genes. Each gene contains at least two variants passing quality control. The first 10 principal components of EIGENSTRAT were included as covariates in the SKAT analysis. We also carried out conditional analyses by including the minor allele count of nearby known common or rare variants as covariates. Additionally we performed a simple burden test to assess if rare variants were enriched in cases versus controls or in controls versus cases. We used the Fisher’s exact statistical framework described for the single-variant association analyses above. To interpret statistical significance, we applied a Bonferroni corrected significance threshold of *P* < 2.06×10^−6^ (*P* = 0.05/12,122 x2 gene-based tests).

Haplotype construction of and frequency calculation of the *CFH* region was performed using the samples in the Boston cohort. Haplotypes were constructed using the --hap-freq function in PLINK and haplotypes with a frequency of greater than 0.001 in either cases or controls were reported (64). Seven SNPs in *CFH* were used in the haplotype construction including the rare *CFH* SNPs, R1210C and N1050Y.

We used the CaTS-Power Calculator software (http://csg.sph.umich.edu//abecasis/cats/calculator.html) to estimate the power to detect each of the rare single-variant associations identified, assuming a significance threshold of *P*-value < 5×10^−08^.

## ACKNOWLEDGEMENTS

We thank the numerous ophthalmologists throughout the United States that contributed to the Boston AMD cohort, the ophthalmologists from the Clinical Researches Functional Unit, CHI Créteil, France, and the ophthalmologists from the Helsinki University Central Hospital, Helsinki, Finland.

## FIGURE LEGENDS

**Supplementary Figure S1. Quantile-Quantile plot of gene-based tests.**

We tested a total of 12,122 genes using the sequence kernel association test (SKAT). We plot the observed *P*-value for each gene as a function of expected *P*-values.

**Supplementary Figure S2. Quantile-Quantile plot of common ancestry informative markers.**

We tested 15,671 ancestry-informative markers excluding regions near (<1Mb) any of the 19 known AMD loci (18), and the major histocompatability complex locus (chr 6, 25-35 MB). We plot the observed *P*-value for each marker as a function of expected *P*-values.

**Supplementary Figure S3. Manhattan Plot of common variants.**

We plot the -logi_10_(*P*-value) for each common variant organized by location on each chromosome.

**Supplementary Figure S4. Quantile-Quantile plot of rare variants.**

We plot the observed *P*-value for each rare variant as a function of expected *P*-values.

**Supplementary Figure S5. Quantile-Quantile plot of common variants for advanced AMD vs unaffected analysis.**

We plot the observed *P*-value in the advanced AMD vs unaffected regression analysis for each common variant as a function of expected **P*-* values.

**Supplementary Figure S6. Quantile-Quantile plot of common variants for neovascular disease vs unaffected analysis.**

We plot the observed *P*-value in the neovascular disease vs unaffected regression analysis for each common variant as a function of expected *P*-values.

**Supplementary Figure S7. Quantile-Quantile plot of common variants for geographic atrophy vs unaffected analysis.**

We plot the observed *P*-value in the geographic atrophy vs unaffected regression analysis for each common variant as a function of expected *P*-values.

**Supplementary Figure S8. Quantile-Quantile plot of common variants for geographic atrophy vs neovascular disease analysis.**

We plot the observed *P*-value in the geographic atrophy vs neovascular disease analysis for each common variant as a function of expected **P*-* values.

**Supplementary Figure S9. Principal component analysis including all cases and control samples.**

We plot the eigenvector1 values for each case and control sample as a function of engenvector2 values to show no stratification between cases and controls is present in the dataset. Cases = red +; controls = green x.

